# AI-based graptolite identification improves shale gas exploration

**DOI:** 10.1101/2022.01.17.476477

**Authors:** Zhi-Bin Niu, Hong-He Xu

**Affiliations:** State Key Laboratory of Palaeobiology and Stratigraphy, Nanjing Institute of Geology and Palaeontology and Centre for Excellence in Life and Paleoenvironment, Chinese Academy of Sciences, 210008 Nanjing, China; College of Intelligence and Computing, Tianjin University, 300354 Tianjin, China

**Author notes:** These authors contributed equally to this work.

## Abstract

Graptolites are fossils from the mid-Cambrian to lower Carboniferous periods that inform both our understanding of evolution and the exploration of shale gas [1–4]. The identification of graptolite species remains a visual task carried out by experienced taxonomists because their fine-grained morphologies and incomplete preservation in multi-fossil samples have hindered automation. Artificial intelligence (AI) holds great promise for transforming such meticulous tasks, and has already proven useful in applications ranging from animal classification to medical diagnostics [5–15]. Here we demonstrate that graptolites can be identified with taxonomist accuracy using a deep learning model [16–18]. We develop a convolutional neural network to classify macrofossils, and construct a comprehensive dataset of >34,000 images of 113 graptolite species annotated at pixel-level resolution to train the model. We validate the model’s performance by comparing its ability to identify 100 images of graptolite species that are significant for rock dating and shale gas exploration with 21 experienced taxonomists from research institutes and the shale gas industry. Our model achieves 86% and 81% accuracy when identifying the genus and species of graptolites, respectively; outperforming taxonomists in terms of accuracy, time, and generalization. By investigating the decisions made by the neural network, we further show that it can recognise fine-grained morphological details better than taxonomists. Our AI approach, providing taxonomist-level graptolite identification, can be deployed on web and mobile apps to extend graptolite identification beyond research institutes and improve the efficiency of shale gas exploration.

---

Accurate and efficient species identification of graptolite, an extinct group of globally distributed and rapidly evolved macrofossil [1, 19], is indispensable in research on evolution and biostratigraphy and assisting global shale gas exploration [3]. There were 102 Ordovician and Silurian graptolite species selected as biozones for determining global rock age and regional correlation, and contributing to understanding the evolutionary pattern of ancient life [19], and 16 graptolite species as “gold callipers” to locate shale gas favourable exploration beds (FEBs) in China [[3], Extended Data Fig. 2]. Graptolite identification has to date been an exclusively human task in palaeontology. We developed a computational approach that decreases palaeontologist workloads and enables shale gas specialists to accurately identify graptolite species and find shale gas FEBs in seconds. By creating a unique dataset of authoritative, taxonomical, and pixel-level annotated graptolite images, we are able to train the world’s first deep neural network based taxonomist-level, macrofossil identification model.

Previously, the identification of graptolite species has mainly relied on human visual inspections of specimens, rather than methods involving chemistry, spectrum, molecular biology, genomics, or histology. Graptolite organisms have lost their soft tissues during fossilization and most are found flattened and carbonized. Palaeontology taxonomists identify graptolite based on examinations using a hand lens in the fieldwork, whilst determine accurate species after removing the coverings and applying diagnostic measurements aided with microscope in the laboratory. The identification requires lengthy specific training based on mass observations of limited specimens that are mainly housed in academic institutes, and consequently, the number of qualified graptolite taxonomists cannot meet the massive and urgent demand required for geologic survey and shale gas exploration [2, 3]. Shale gas companies have had to deliver the specimens to research institutes for accurate identification, which is a time-consuming and costly procedure.

Although computer-aided fossil identification (CAFI) software was introduced as early as 1980s [12] with several machine learning based approaches have been proposed hereafter [20–23], to our knowledge, no CAFI assistance tool has been used for oil/gas exploration due to the lack of generalization capabilities, primarily caused by insufficient labelled fossil images followed by limited performance of the shallow machine learning algorithms. It was until recent years, the deep learning methodology reformed the prediction framework, showed promising results in medical diagnoses [5–7, 24] and many other fields, illuminated a possibility of industrial level CAFI. Identification of microfossils, such as foraminifer[8], pollen grains [9–11], and micro-objects from thin sections [12–14], saw a sprout of interdisciplinary trend of palaeontology and artificial intelligence[25, 26]. Microfossils, before photographing, are primarily treated by dissolving the impurities and surrounding rocks, so they are relatively intact, isolated, uniform textured, and rarely overlay, and can be massively photographed cost-effectively using automated slide scanners. The automated identification of macrofossils, e.g., animal bone and plant leaf fossils, is more challenging, because they are often with incomplete preservation, clutter and occlusion of multi-fossil remains, a lack of contracts between the fossil remains and surrounding rocks, colour confusion caused by mineralization, and varying developmental states. Besides, their specimens and images are more difficult to obtain.

We fill the gap by creating a unique dataset of annotated macrofossil graptolite images and training a state-of-the-art deep learning model (Fig. 1). The existing body of work are either mainly conducted on small scale dataset or focus on microfossils [8, 9, 11, 21, 27], thus does not generalize well to macrofossils, the more challenging direct evidence of evolution. The limited automated macrofossil identification has usually required formatted traits[12] with landmarks, and even manual measurements; by contrast, our system requires no hand-crafted features only pixels. We demonstrate generalizable classification ability with the first macrofossil dataset of 34,620 pixel-level annotated graptolite images. We highlight the development of the AI approach and its ability to outperform graptolite taxonomists in image-based identification in terms of accuracy, time cost, and generalization capability.

**Fig. 1.**
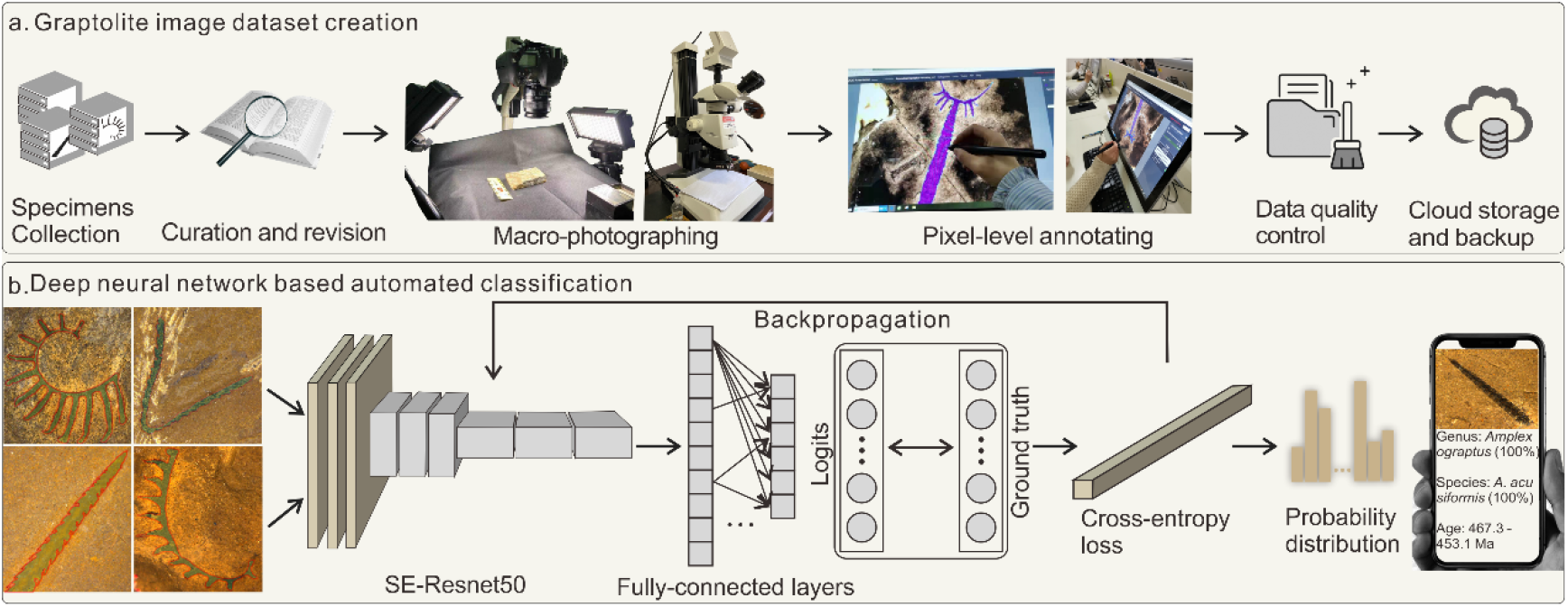
The AI approach workflow. **a**, The process of creating the dataset. The graptolite specimens were carefully curated and revised to select the species with biostratigraphy and application significance. Every image was obtained from specimens that were macro-photographed using a single-lens reflex camera and microscope. They were then pixel-level annotated according to the graptolite body’s contours. After professional revision and cleaning of the annotated images, the whole dataset was uploaded to and stored in our cloud server. **b**, The process of the supervised deep neural network based automated classification. We evaluated nine state-of-the-art CNN models for the dataset (see Extended Data, Fig. 4, Table 1) and chose the squeeze-and-excitation network built on Resnet-50 [17] as our main network owing to its superior performance with our dataset. The offline trained model can be deployed on the cloud server to support remote identification from a web interface or mobile device. Notably, the end users are required to annotate the contours of the fossil body for more accurate identification.

## Pixel-level annotated graptolite image dataset

Our dataset consists of 1,565 pieces of specimens collected from 154 representative geological sections of China (Fig. 2). These specimens were carefully curated, and taxonomically belonging to 113 graptolite species or subspecies, of 41 genera, 16 families of the Order Graptoloidea (Extended Data, Fig. 2, Appendix 1). We incorporated revision suggestions of several distinguished palaeontologists were incorporated to generate the ground-truth labels for the supervised training, providing a taxonomical authority of the dataset. These graptolite species are commonly used in geological age determination and shale gas FEB indication, including 22 graptolite biozones from the Dapingian stage of the Ordovician (470.0 Ma) to the Homerian stage of the Silurian (427.4 Ma), and 16 “gold callipers” of shale gas FEBs for cases of 20 to 80 m thick graptolite shale in China [3, 28] (Fig. 2, Extended Data, Fig.2).

**Fig. 2.**
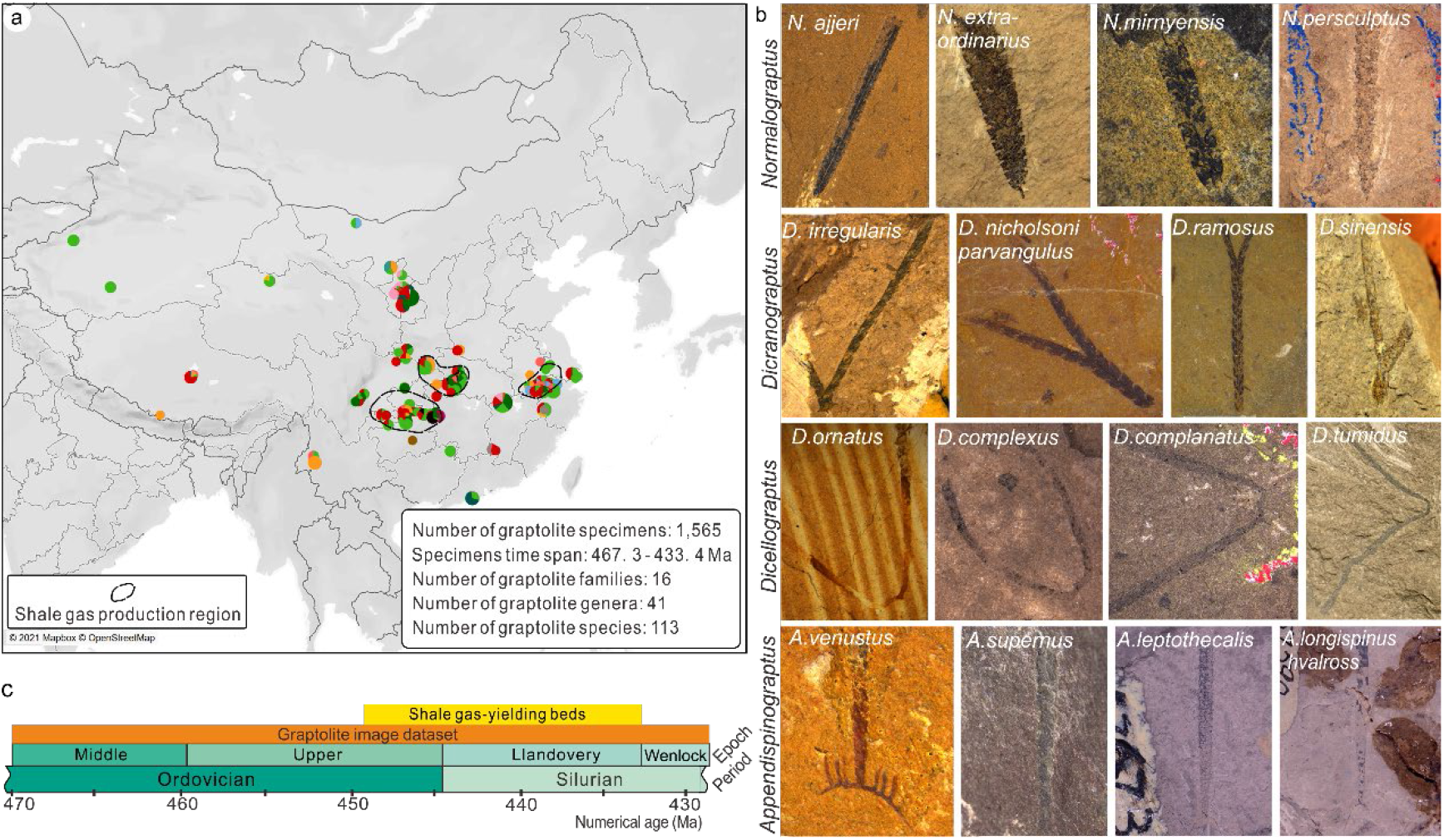
Geological significance and example graptolite images of the dataset. **a**, The graptolite specimens’ geographic distribution and statistical results of our dataset. Each locality is represented by a pie chart where each colour is encoded as one graptolite family of the order Graptoloidea. The sector size is proportional to the specimen number for every family. The radius of the pie chart is proportional to the total number of specimens from the same locality. The dashed-lines circle the main areas of shale gas production. **b**, Example graptolite images. Each row represents a same genus with different species graptolites showing the high intra-class and low inter-class variance nature of the graptolites. **c**, These graptolite species range from the Middle Ordovician to the Wenlock of the Silurian (black line section), covering all shale gas-yielding beds from southern China (yellow bar).

We photographed every specimen under different focusing angles and scales and obtained a total of 40,597 images. The image dataset was cleaned and professionally revised. We removed unclear images and those with questionable taxonomical statuses. Every image was pixel-level annotated, according to the graptolite’s body contours (Fig. 1a). Ultimately, our dataset contains 37,588 images for training, validating, and testing (Extended Data, Fig. 3, Appendix 1).

## Deep neural network-based automated taxonomical classification

We chose to use Convolutional Neural Network (CNN) as the supervised learning classifier for the AI based automated taxonomical classification. We conducted experiments and compared the performance of 9 state-of-the-art CNN models on the graptolite image dataset (Extended Data Table 1, Fig.4), and selected to use the squeeze-and-excitation networks [29, 30] built on Resnet-50 (SE-Resnet50) as the AI engine for its optimal performance on our dataset.

The SE-Resnet50 architecture uses a transfer learning manner to train the model (see Fig. 1b). The using of pre-trained weights leveraging the natural image features learned by ImageNet [31, 32] accelerated convergence, especially in the early stages of training. We then fine-tuned the parameters across all layers of our dataset. During the training, we used the stochastic gradient descent optimizer, with a momentum of 0.9 and weight decay of 5 × 10^−4^. The initial learning rate was set to 0.001 and decreased following a cosine annealing schedule. It slowly reduced to approximately 2.7 × 10^−8^ in the last epoch. We used the cross-entropy loss function to calculate the model error. As in Figure 1, the gradients were computed using backpropagation of the error algorithm and the parameters updated using the stochastic gradient descent optimization algorithm. After training each epoch, we fed all test images into the model, performed forward propagation to calculate accuracy, and saved the model parameters with the highest accuracy as our final training model.

## Cross-validation of genus- and species-level identification

We first conducted a cross-validation to evaluate the hyper-parameters and generalizability. As the number of specimens of each species ranged from 4 to 20 (Appendix 1), we chose to use three-fold cross-validation to evaluate the performance, so that the cross-validation would cover all species. The validation image set was carefully chosen so that every specimen-related image was completely removed from the training set, and thus there was no intersection between the training and validation test sets. The validation set included 15 to 80 randomly chosen images of each species. In total, there were 34,620 images for training and validation; each iteration had a different number in the validation set (Extended Data, Table 1). We conducted two cross-validations on different taxonomical levels. On the genus and species levels, the AI-based approach achieved 81.8% and 70.538% overall accuracy, respectively (Extended Data, Table 1). These results indicate that the AI approach is a feasible means of learning the ultra-fine-grained features of graptolites.

## Comparison of the AI-based approach and graptolite taxonomists

The cross-validation results were promising but still inconclusive, as the results depended on a particular random choice for the pairing of training and validation sample sets. To test the generalization ability of the model and conclusively evaluate our AI approach, we further compared the performances of our model with graptolite taxonomists on a carefully chosen “golden standard” test set.

We chose 100 images of 35 graptolite species to conduct the golden standard test. These species are directly related to dating sediments and locating shale gas FEBs during the mining. They consist of all 22 graptolite biozone species from the Middle Ordovician to the Wenlock of Silurian and 16 indicator species widely used in shale gas exploration in China (Extended Data, Fig.2). 35 species of the golden standard test set are from 99 pieces of specimens, which are dubbed as “collection-99” specimen dataset. There are totally 2,968 photographs taken from the collection-99, 100 images selected into the golden standard test set are well-focused, and friendly showing morphological characters of graptolite for identification. The rest of images taken from the collection-99, 2,868 images, are removed from the training set. Finally, we use 31,652 images to train the AI model.

We have invited 21 graptolite taxonomists to provide a baseline of human performance. All experts are working on graptolite-related education, research, and engineering applications at universities, institutes, or oil/gas companies around the world. They have 5 to 30 years of fossil (especially graptolite) identification experience. During the golden standard test, experts were asked to identify the graptolite and record the genus and species names, time cost, and comments (Appendix 4). They could view and refer any literatures even internet searching freely.

We use identification accuracy, time cost, and micro-average sensitivity-specificity curves (i.e., receiver operating characteristics curves, ROC) to evaluate the performances of the AI approach and human’s classification ability (Fig. 3). Our AI approach achieves 86% and 81% levels of accuracy for genus and species identification respectively in seconds, whilst those of the graptolite taxonomists are only 59.5% (95%CI 52.6%, 66.5%), and 45.2% (95%CI 38.9%, 51.5%), taking on average 4.8 minutes (95%CI 4.2 min, 5.4 min) (Extended Data, Table 2). For further details, in genus identification, the academic experts achieve 68.0% (95%CI 60.0%, 73.5%) and industry experts achieve 45.3% (95CI 33.0%, 57.6%); while in species, the numbers are 50.8% (95CI 44.7%, 56.8%) and 35.8% (95CI 21.0%, 50.6%), respectively (Fig. 3a).

**Fig. 3.**
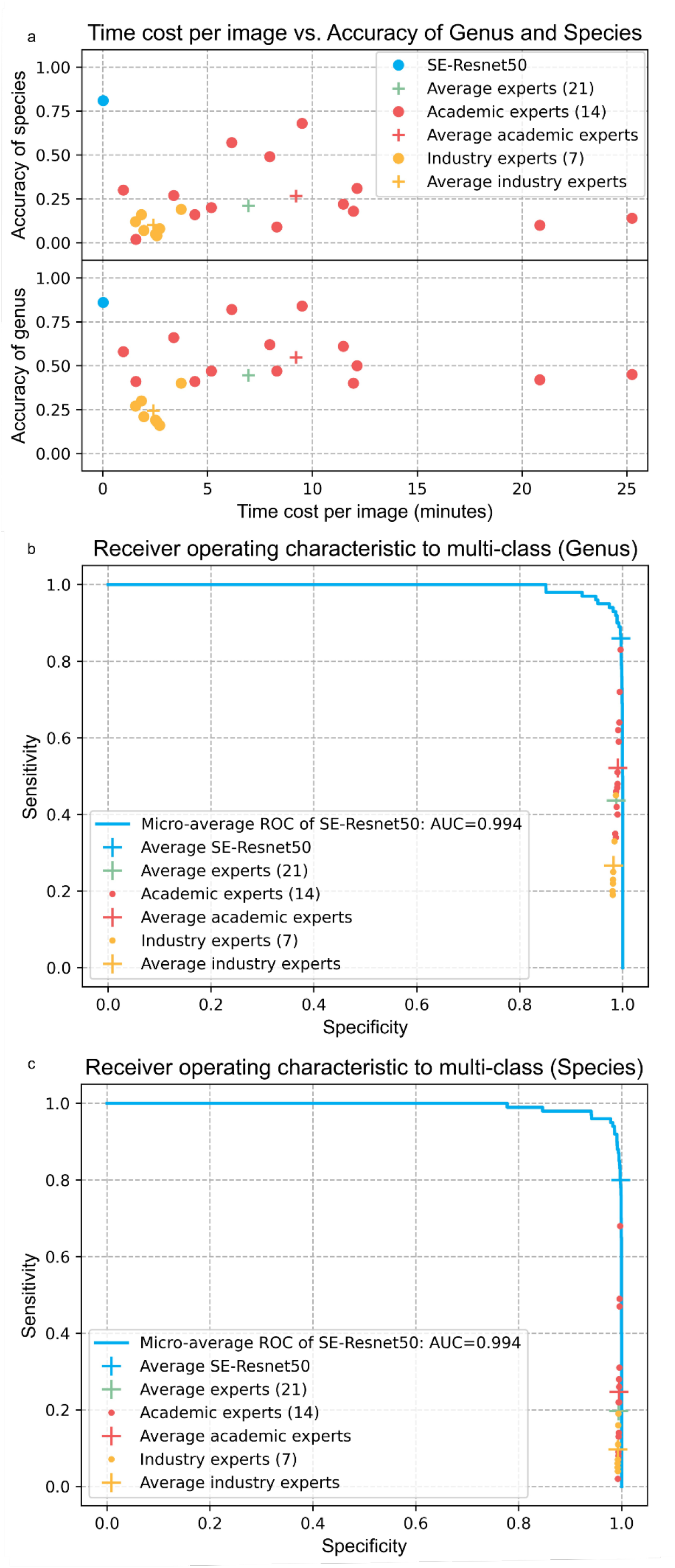
Performances of AI approach and experts on gold standard graptolite identification test. a, Time cost - genus and species identification accuracy plot. It shows that the AI based approach (SE-Resnet50) significantly outperforms the experts in genus and species graptolite identification accuracies and the efficiency. The average accuracy of academic experts is higher than that of industry experts at both two levels, indicating that the academic experts can make more reliable predictions, but they usually take longer time. b, Comparison of the micro-average roc curve of SE-Resnet50 with the micro-average experts at genus level. The average point of SE-Resnet50 is located at the upper than that of all experts, indicating that the classification performance of the model is evidently better than that of all experts. The average point of academic experts is located at the upper right of that of industry experts, which means that academic experts have better classification ability. b, The comparison of the micro-average roc curve of SE-Resnet50 with the micro-average experts at species level. The average point of SE-Resnet50 is located at the upper right of all experts, which indicates that the classification performance of the model is obviously better than that of all experts. The average point of academic experts is located at the upper right of that of industry experts, which means that academic experts have better classification ability.

**Fig. 4.**
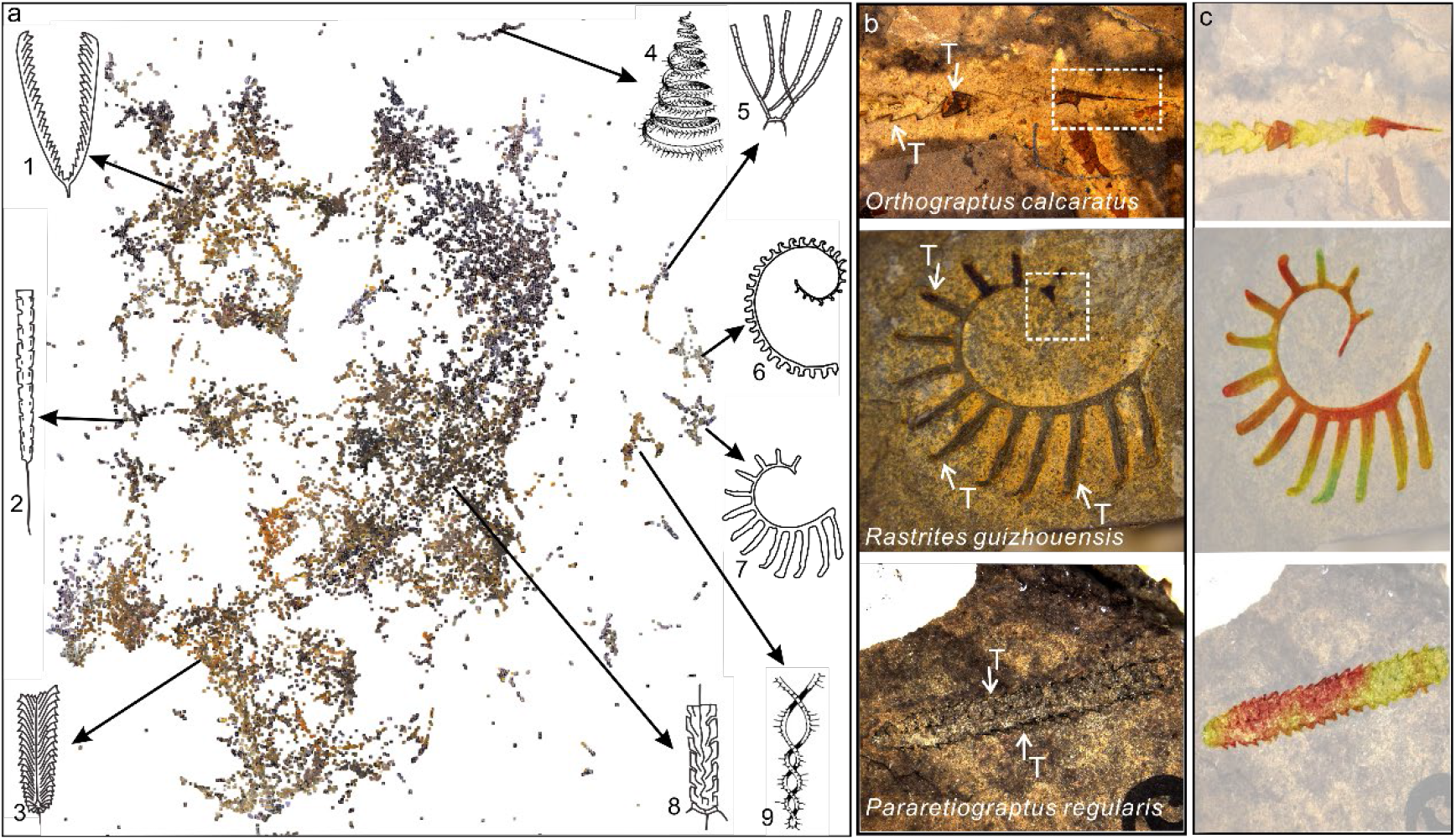
Visualisation of our graptolite image dataset structure and attention map of example graptolite images. **a.** UMAP embeddings of the last hidden layer representations of our dataset. Every image is projected to a thumbnail on the two-dimensional scatterplot, visually revealing at least nine graptolite morphotype clusters (Extended Data Table 3). **b**, 3 example images from the gold standard test set, clearly showing graptolite thecae (short or long prominence structures, arrowed, T) and proximal ends (dashed box ) of 3 species, providing good and sufficient morphological information for expert’s identification. No scale bar is given in every image and it is impossible to measure fossil remains, such might affect the species identification of some graptolite. The proximal end is now visible in the lowermost graptolite specimen image, making identification impossible to experts but feasible to the AI model. **c**, 3 images with highlighted discriminative visual regions generated through CAM. We use a rainbow colour map to reweight and render the saliency regions used for image classification. Red regions correspond to high score for class, the next are yellow and green. The high score regions of the AI model coincide with critical areas of expert’s identification, e.g. the proximal end, whilst to the graptolite without showing the proximal end, the AI model recognizes regular thecae as visual pattern and better identify the graptolite than experts.

We provide micro-average sensitivity-specificity curves to quantitatively measure the classification performances of the AI model and experts (Fig.3bc, Method). In genus identification, the experts achieve on average sensitivities of 43.67% (95%CI 35.7%,51.7%), specificity of 98.8% (95%CI 98.6%, 98.9%), and in species level, 19.7% (95%CI 11.9%, 27.5%) and 99.4% (95%CI 99.3%, 99.5%) respectively. The AI model has 86% sensitivity and 99.7% specificity (genus level), and 80% sensitivity and 99.8% specificity (species level). Overall, the AI model outperforms the experts in genus 0.89% (95CI 0.68%, 1.1%) specificity and 42.3% (95%CI 34.4%, 50.3%) sensitivity; in species 0.40% (95CI 0.34%, 0.47%) specificity and 60.3% (95%CI 52.5%, 68.1%) sensitivity.

The AUC (Area under the ROC Curve) illustrates how much the model is capable of distinguishing between classes. The micro-average AUC value is near 1.0 (99.4%), indicating that the AI model is generally able to precisely identify the graptolite taxonomy. The human-machine contest suggests that our AI approach is significantly superior to over-5-years-experenced academic graptolite taxonomists and industry specialists, thus therefore, the relative expensive and time-consuming graptolite identification in the fieldwork would benefit more from the potential CAFI software.

## Interpretation through visualisation of the AI graptolite identification internals

We examined the features learned by the deep neural network through uniform manifold approximation and projection (UMAP) [27] (Fig. 4a). Each thumbnail image in the two-dimensional space represents a graptolite projected from the 100,352 dimensional output of the CNN’s last hidden layer [28]. The proximity in low dimensional UMAP space reveals nine major graptolite morphotype clusters [33] (Fig. 4a, line drawings showing the major morphotypes of the clusters). In the middle of the map is the dominant cluster, namely the scandent morphotype, including almost all biserial diplograptids. The upper left portion of the map is the V-shaped graptolite morphotype, including all graptolites with typical U- or V-shaped tubariums. We also found that several outliers on the right part of the embedding had distinctive morphology, such as graptolite with spiral or helical shapes (Fig. 4a, Extended Data Table 4).

We further utilized class activation mapping (CAM) [26, 35] to visualise and interpret the prediction decisions made by the deep neural network (Fig. 4c). The attention map indicates the diagnostic (semantic) image regions by projecting back the weights of the output layer on to the convolutional feature maps, localizing class-discriminative regions that influence the AI-based approach and showing the differences between expert and the AI model dealing with graptolite images. The morphology of proximal end (sicula and one or two thecae, the area in the dashed line box in Fig. 4b) is critical to identify graptolite genus and superior genus taxa, and measurements are necessary to identify some graptolite species, especially within the genus [1]. Actually, proximal ends are not either readily preserved in every specimen or fully given in our test set images, to which, as a consequence, identification is impossible to experts, except graptolite of specialized type, as shown in the third line of Fig. 4b. Additionally, the AI model recognizes visual patterns of images as important class-discriminative information better than experts. For example, graptolite with only regular thecae is also identifiable to the AI model but not to experts. The attention map demonstrates that the superhuman performance of the AI approach is able to recognize morphological nuances of graptolite species without requiring measurements, comprehensively incorporating ultra-fine grained details, including “traits agnostic” of graptolite.

## Discussion

We demonstrate the effectiveness of deep learning in macrofossil graptolite identification, an AI-based approach applicable to species-level graptolite identification; the approach exhibits supertaxonmist performance, sufficient generalization, and interpretable semantics. Graptolite shale comprises over 9% of the hydrocarbon rocks in the world (Extended Data, Fig. 1) and yields over 61.4% natural gas for China. This AI approach potentially reshapes the macrofossil identification and the outfitted with smart mobile device can improve greatly the efficiencies of geological surveys and shale gas exploration. This research contributes to the transdisciplinary future of palaeontology, artificial intelligence, and the oil/gas industry. This method is primarily constrained by limited data sources in China. Further work will incorporate much wider or perhaps global sourced data. Future research may focus on fossils from more broad groups, integrating expert’s experience and knowledge to the AI-based approach.

## Methods

### Dataset

All graptolite images were taken from 1,565 pieces of specimens (Appendix 1), which are housed at the Nanjing Institute of Geology and Palaeontology (NIGP), Chinese Academy of Sciences (CAS) – the world largest palaeontological research centre and one of the top 3 specimen collection centres. The NIGP boasts approximately 180 palaeontological researchers and laboratory technicians and collecting over pieces of fossil specimens [35]. All specimens were professionally curated and photographed using single-lens reflex digital Nikon D800E cameras with Nikkor 60 mm macro-lens and Leica M125 and M205C microscopes equipped with Leica cameras.

We hired 7 data entry clerks and 3 engineers were hired for specimen photography, pixel-level annotation, and data cleaning. We spent over 2 years to complete the dataset. In total, we took 40,597 images, including 20,644 camera photos (each with a resolution of 4,912 × 7,360) and 19,953 microscope photos (each with a resolution of 2,720 × 2,048). The entire volume of the dataset is 299 GB. We examined all photos, removed 5,977 unclear images or those with questionable taxonomical status, and keep 34,620 valid images in our dataset for validation, training, and testing. We employed three-fold validation; In each iteration, 2,932 to 3,051 images were used for validation and the rest for the training. After the validation, the dataset is separated into a train set of 31,652 images and a golden standard test set of 100 images, which contains 22 photos taken using single lens reflect camera and 78 with microscope, were photographed from 99 pieces of specimens.

### Data Preparation

The data cleaning was conducted by graptolite palaeontologists from the NIGP at the CAS. The graptolite images were removed if they were: 1) incorrect or poorly focused, with very low contrast; 2) of poorly preserved specimens and substantially deformed; 3) incorrectly labelled and impossible to identify; or 4) showing graptolite bodies with confusing textural information that hindered the ability to see the morphological characteristics. During the data cleaning process or poorly focused images were removed from the test and validation sets but were still used for training. Images showing graptolite that was partially destroyed or with lesions remained in the dataset. These images were used in training, but extensive care was taken to ensure that they were not split between the training and validation sets. Our data entry team annotated the graptolite bodies at a pixel-level using COCO annotator [36], a web-based image annotation tool. The images were then stored and backed up in the cloud server.

### Training procedure and settings

The training images were initially processed through an augmentation and normalization operation. In detail, the pixel-level annotated graptolite images were resized to 480 × 480 pixels by up-sampling or down-sampling, through interpolation and paddling each graptolite body’s surrounding area with zero background. To the utmost extent possible, the graptolite body aspect ratio in each image was kept constant. The images were then randomly cropped to 448 × 448 pixels. This served as the input size for the deep neural network and was an empirical trade-off between learning accuracy and information redundancy in terms of fine-grained image classification [29, 37–40]. We further augmented the images by flipping, rotating, and colour jittering operations in order to enhance the model’s generalize ability (e.g., to enable the model to adapt to various test images). The images were then normalized with the Z-score algorithm on each image channel. This operation further accelerated the optimization convergence speed. Similarly, the test graptolite images were also centre-cropped and normalized.

We choose to use Convolutional Neural Network (CNN) as the supervised learning classifiers for the AI based automated taxonomical classification. We conduct experiments and compared the performance of 9 state-of-the-art CNN models on the graptolite image dataset (Extended Data Table 1, Fig.4), and choose to use the squeeze-and-excitation networks [29, 30] built on Resnet-50 (SE-Resnet50) for its best performance on our dataset as a representative AI based approach. We give details of the SE-Resnet50 architecture here. The deep neural network was pretrained on approximately 1.3 million images of the ImageNet dataset[31, 32]. The pre-trained model was believed to extract sufficient general image information for feature extraction [5, 41]. In practice, we copied the parameters of the pre-trained model, and fine-tuned them based on the target data set, achieving better results than from the random initialization. The use of pre-trained weights can accelerate convergence, especially in the early stages of training. We then used out dataset to fine-tune the parameters across all layers.

We trained the model for 300 epochs with a mini-batch of 32. In each epoch, we first shuffled all images in training set, and then iteratively fed 32 mini-batch images into the model until all the training images were loaded. The convolutional and pooling layers turned them into a 32 × 2048 × 1 × 1 feature map. Each feature map was flattened to a 32 × 2048 matrix and then mapped to the label space through the fully connected layer. Lastly, a softmax layer was employed to obtain the probability distribution matrix of 32 × 113, giving the probability of each image belonging to each of the 113 taxonomies. During training, we used a stochastic gradient descent optimizer with a momentum of 0.9 and weight decay of 5 ×10−4. The initial learning rate was set to 0.001 and decreased following a cosine annealing schedule. Thus, the learning rate initially reduced slowly, then accelerated, and then slowly reduced again to about 2.7 × 10−8 in the last epoch. This decay strategy was well combined with the stochastic gradient descent algorithm to optimize the objective function. The learning rate was usually higher in the early stages of training, so the network converged quickly. The learning rate was then reduced to make sure the network better converged to the optimal solution.

In general, neural network learning is an optimization procedure, and we used the cross-entropy loss function to calculate the model error. The problem of learning was cast as a search optimization problem. In our approach, the gradients were computed using the backpropagation of the error algorithm, and the parameters were updated using the stochastic gradient descent optimization algorithm. After training each epoch, we fed all test images into the model and perform forward propagation to calculate accuracy; we saved the model parameters with the highest accuracy as our final training model.

### Sensitivity-specificity curve

The sensitivity-specificity curve is a probability curve at various threshold settings. It is also known as receiver operating characteristic curve, or ROC curve. The area under the curve is named as AUC and represents the degree or measure of separability. Our task is multiclass classification problem; thus, the comparison metrics are defined as macro-average and micro-average sensitivity and specificity:

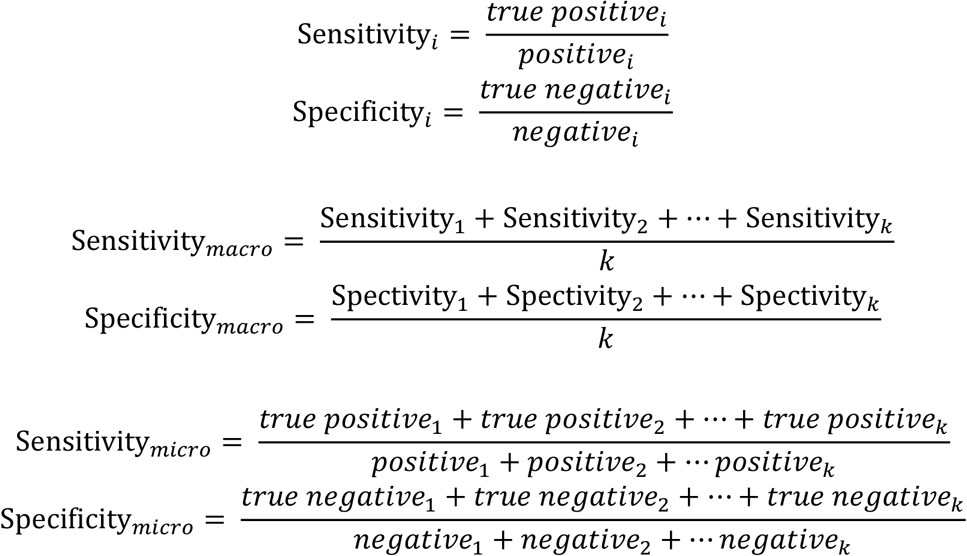

where *true positive_i_* is the number of correctly predicted fossils, *positive_i_* is the number of the type *i* fossil shown, *true negative_i_* is the number of correctly predicted fossils *i*, and *negative* is the number of fossils of type *i* shown. When a test set is fed through the deep network, it outputs a probability, *P*, of fossil types by image. In macro-averaging, all classes are equally weighted when contributing their portion of the value to the total, while in micro-averaging, each observation receives equal weight. This gives greater power to the classes with the most observations. We computed both averaged sensitivity-specificity values by adjusting the threshold of the deep neural network classifier. The sensitivity-specificity curve is a probability curve at various threshold settings and its area under the curve (AUC) represents the degree or measure of separability.

We chose to use micro-instead of macro-averaging because the former computes the metric for each class independently and then averages them, whereas the latter treats all classes equally, and thus therefore, micro-averaging is preferable in multi-class classification tasks with imbalanced classes.

### Confusion matrix

This is a metric for evaluating classification accuracy. Each row of the matrix represents the instances in a predicted class, while each column represents the instances in an actual class (or vice versa). The extended data in Figure 3 provide the confusion matrix for the test set using the AI-based approach.

### Uniform manifold approximation and projection

The uniform manifold approximation and projection (UMAP) algorithm [42], a dimension reduction and visualisation technique, was employed to validate the feature extraction ability of the deep learning model and visualize the distribution of all images. In detail, we reshaped the last convolutional block of the AI model, generating feature maps (2,048 × 7 × 7) into 1 × 100,352 dimensional features and fed them into the open-source tool “pix-plot” to build the UMAP layout. The minimum distance between points in the embedding (parameter “min_distance”) was set to 0.001, and the trade-off between local and global clusters (“n_neighbors”) was set to 6. The “metric” parameter was set to “correlation.”

### Data availability statement

The graptolite images dataset is open accessed at http://www.geobiodiversity.com/

## Code availability

The code for COCO annotator is available at https://github.com/jsbroks/coco-annotator. This annotation tool provides us with objects labelling and polygons annotation functions and returns annotation data in COCO format. The UMAP Visualisation is available at https://github.com/YaleDHLab/pix-plot. This tool can project the image features learned from the model into a two-dimensional space so that the images with similar features can be clustered together. The deep learning framework, Pytorch, is available at https://www.pytorch.org/. This is a python machine learning library based on Torch, which provides functions such as powerful GPU accelerated tensor calculation and deep neural network with automatic derivation. Scripts of our model are available from the corresponding author upon request.

## Acknowledgments

We thank Prof. Renbin Zhan, Prof. Yuandong Zhang, Prof. Xunlai Yuan, Prof. Maoyan Zhu, Prof. Yi Wang, Prof. Bing Huang, Prof. Bo Wang, Dr. Qing Chen, Dr. Lixia Li, and Dr. Xuan Ma Nanjing Institute of Geology and Palaeontology (NIGP), Chinese Academy of Sciences (CAS); Dr. Lucy Muir, National museum of Wales (UK); Dr. Petr Kraft, Charles University (Czech); Dr. Petr Štorch, Institute of Geology of the Czech Academy of Sciences (Czech); David Loydell, University of Portsmouth (UK); Dr. Wenhui Wang, Central South University (China); Dr. Yanyan Song, University of Chinese Academy of Sciences; Dr. Ming Li, Chinese Academy of Geosciences; Drs Xin Wang and Jian Wang, Xi ’an Geological Survey Centre, China Geological Survey; Hongyan Wang, Feng Liang, Qun Zhao, Shasha Sun, National Energy Shale Gas R & D (Experiment) Centre, Beijing, China; Mr. Jie Tian, China Petroleum Pipeline Bureau, Langfang, China; Prof. Min Zhu and Dr. Zhaohui Pan, Institute of Vertebrate Paleontology and Paleoanthropology, CAS; and a number of anonymous experts, for comments and suggestions; Prof. Wenwei Yuan and Daojun Yuan, NIGP, CAS, for the courtesy in graptolite specimens examination; Mr. Zhiqiu Zhang, Tianjin University, for testing the software; Xiaojing Tong, Yutong Sun, Xiaoyan Dong, Shuangshuang Song, Danni Fan, Yanyan Zhu, and Qing Xia, NIGP, CAS, for preparations of specimens image data; Yansen Chen, NIGPAS, CAS, for technician supports.

We would also like to thank Dr. Lesley Anson for the constructive comments and suggestions on expression.

This research was supported by the Strategic Priority Research Program of Chinese Academy of Sciences (Grant Nos: XDA19050101, XDB26000000) and the National Natural Science Foundation of China (Grant Nos.: 41772012, 61802278).

## Author Contributions

Z. Niu and H. Xu conceived of the whole project. H. Xu and Z. Niu wrote and discussed the manuscript. Z. Niu and Y Pan realized the code and software. H. Xu, Z. Niu and Y. Pan conducted the experimental analyses.

---

**Extended Data Fig. 1.**
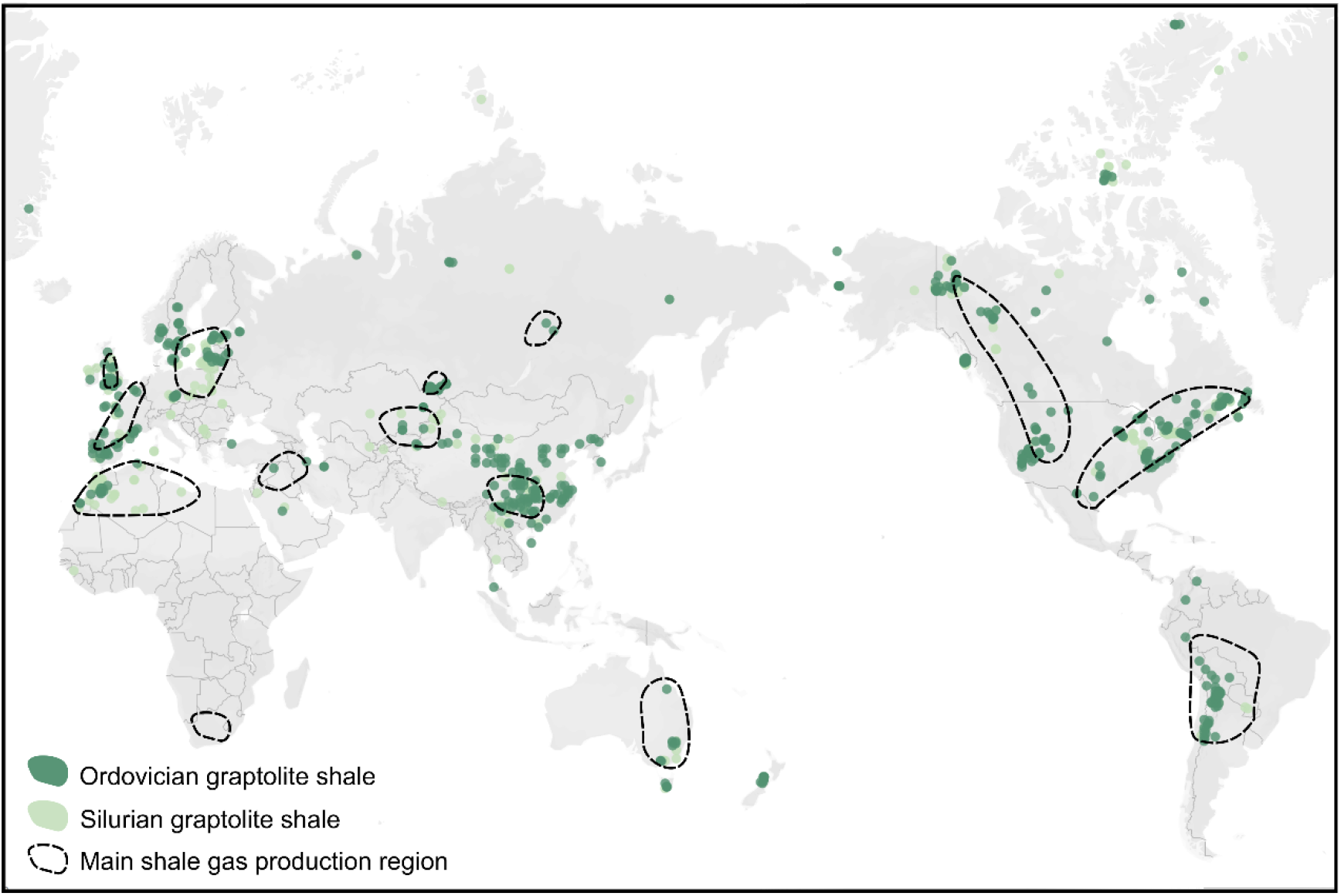
Global distribution of graptolite shale and shale gas regions. Most graptolites are yielded from shales and their distribution is based on graptolite fossil occurrence records in global Ordovician and Silurian sediments [43] (PBDB, http://www.paleobiodb.org/; GBDB, http://www.geobiodiversity.com/). Graptolite shale comprises more than 9% of hydrocarbons rocks and yields the largest volume of shale gas in the world [4, 44]. In China, over 61.4% of the natural gas is yielded from the Ordovician and Silurian graptolite shale of southern China [3].

**Extended Data Fig. 2.**
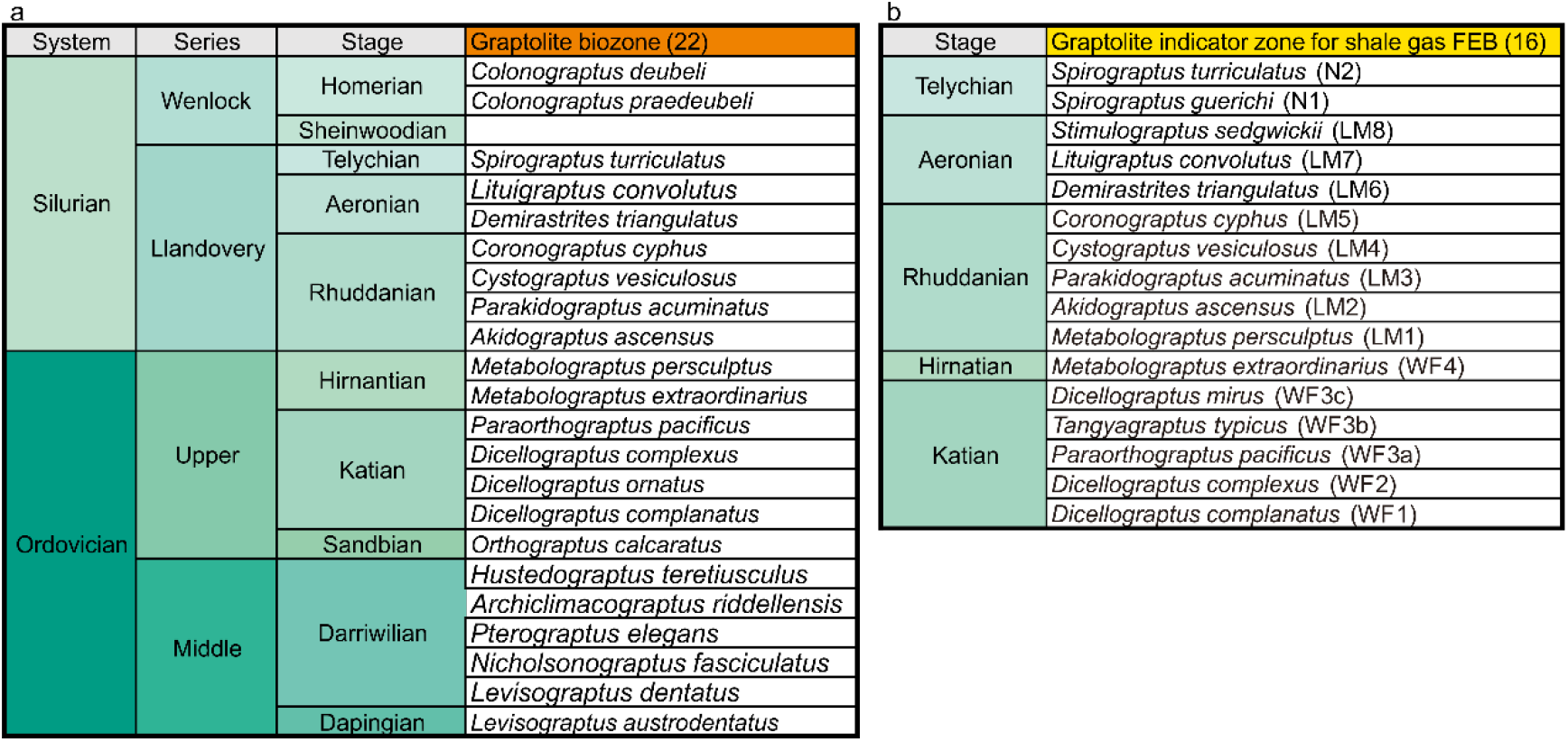
Graptolite species selected as global biozone and indicator zone for shale gas favourable exploration beds of our data set. Graptolite is marine colonial organic-walled hemichordate and has over 210 genera/3,000 species fossil records extending from the Cambrian to the Carboniferous (c. 510~320 Ma). Among our dataset of 113 graptolite species, there are 35 graptolite species to conduct the golden standard test, including 22 graptolite biozones from the Middle Ordovician to (470.0 Ma) to the Wenlock of Silurian period (427.4 Ma) [19], and 16 graptolite species as ‘gold calliper’ to locate favourable exploration beds (FEB) of shale gas in China [3]. Note that some graptolite species are duplicate in the two lists.

**Extended Data Fig. 3.**
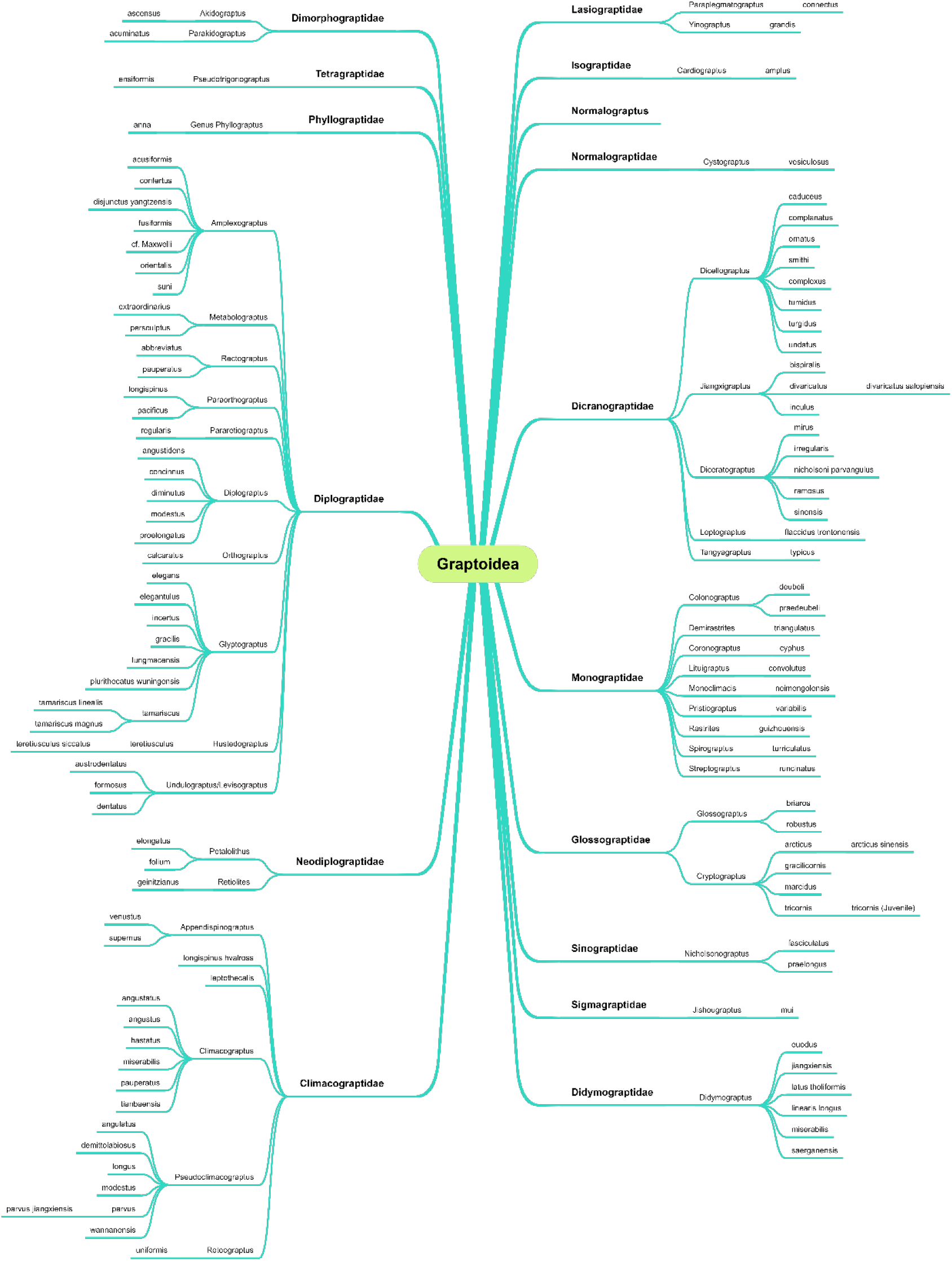
Taxonomy of the graptolites of our dataset. Our dataset consists of 113 species or subspecies, 41 genera, 16 families of the Order Graptoloidea [1]. Revisions and detailed taxa are given in Appendix 1

**Extended Data Fig. 4.**
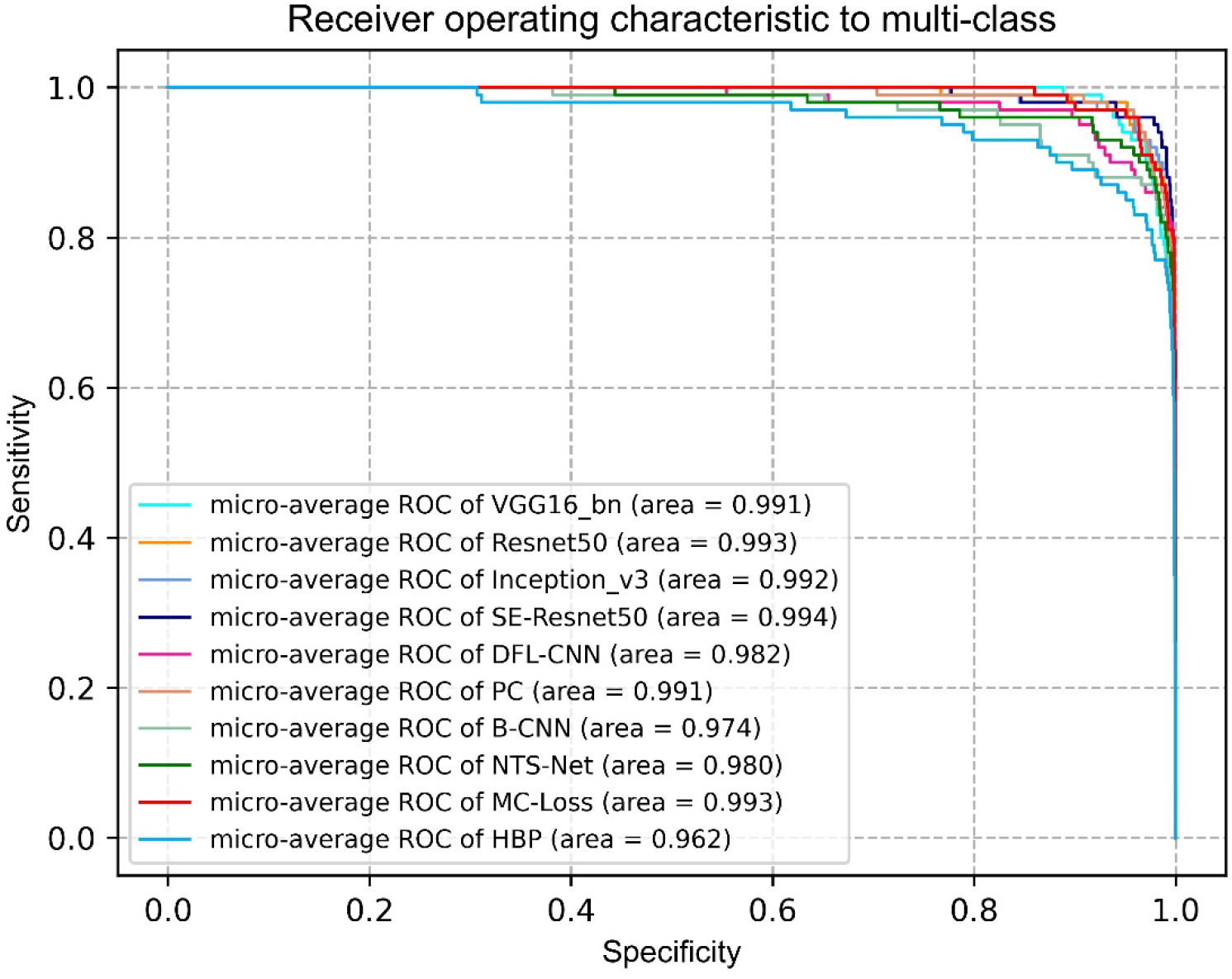
Performance comparisons of nine state-of-the-art CNN models (Extended Data Table 1) for the gold standard graptolite image test set through micro-average specificity-sensitivity curves. The micro-average ROCs give a relative objective quantified of the performance of the classifiers. All these CNN models achieve a preferred performance with an AUC of 98.4% (95%CI 97.4%,99.3%); whereas, the SE-Resnet50 achieves the best performance (99.4% AUC). In the chart, the * means the network uses Resnet50 as backbone and # refers the networks uses VGG16 as backbone. Notably, the vanilla neural networks perform better than the models based on them with our dataset, probably because most of the improved algorithms are designed for large objects with obvious structural information such as birds and planes and does not generalize well on the objects with a single main structure. Based on the conclusion of the comparisons, we choose to use the SE-Resnet50 as a baseline of the AI based graptolite identification.

**Extended Data Fig. 5.**
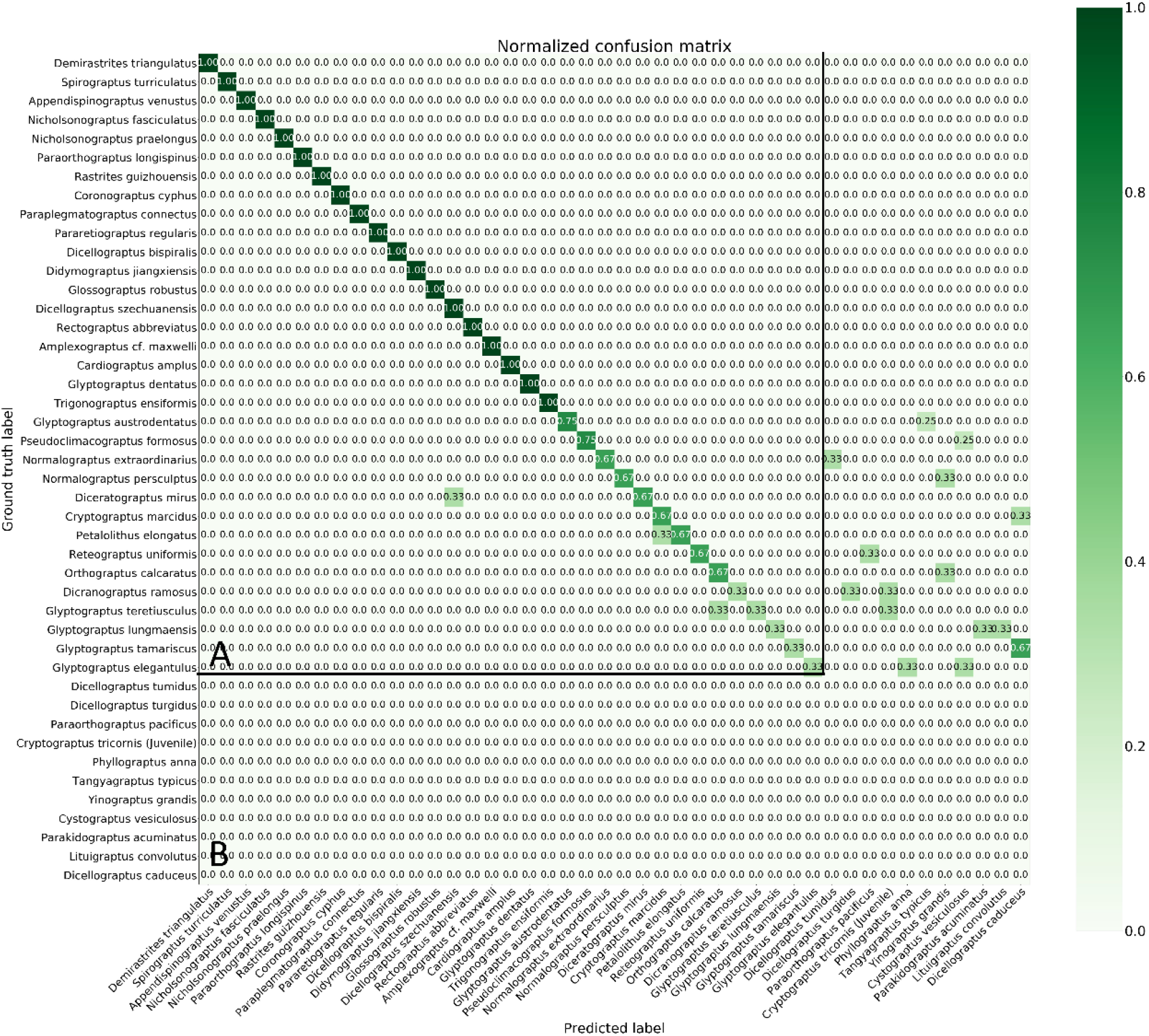
Augmented confusion table of the golden standard test on species classification. Each row is the ground truth label of the image, and the column is the predicted label. The ground-truth label belongs to 35 species or subspecies of graptolite (area A), while some of the images are predicted out of the range (area B).

**Extended Data Table 1.**
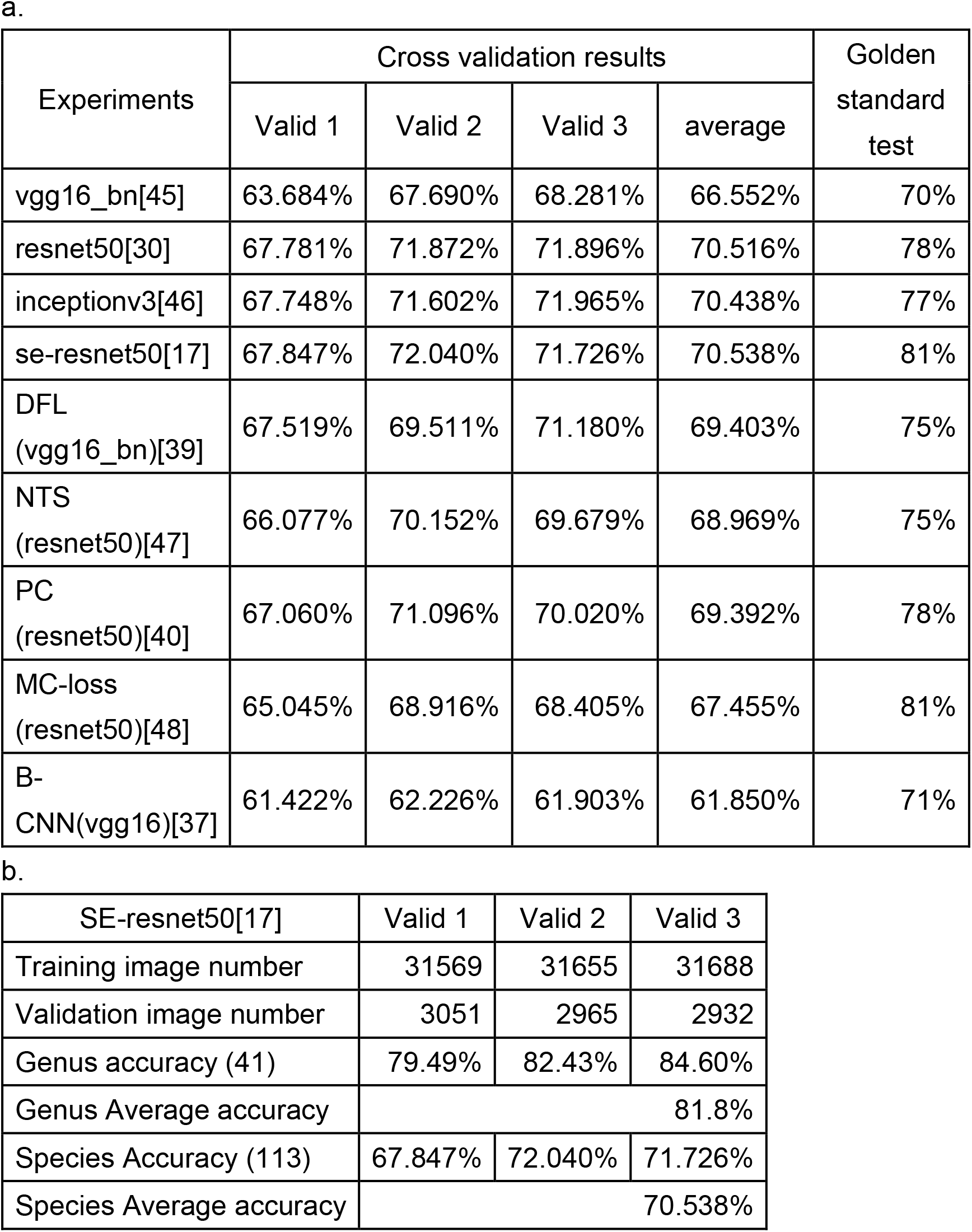
Performance evaluation results. **a**. Three-fold cross validation and test set accuracy using the nine CNN models. In the three fold cross validation, we have 34620 images, with the three fold training and validation image numbers are (31569, 3051), (31655, 2965), (31688, 2932). We observe the SE-Resnet50 gives the best performance. We also give golden standard test on the “testset-99” in the table. **b**. Cross validation for genus and species identification accuracy using se-resnet50.

| Experiments | Cross validation results |  |  |  | Golden standard test |
| --- | --- | --- | --- | --- | --- |
|  | Valid 1 | Valid 2 | Valid 3 | average |  |
| vgg16_bn[45] | 63.684% | 67.690% | 68.281% | 66.552% | 70% |
| resnet50[30] | 67.781% | 71.872% | 71.896% | 70.516% | 78% |
| inceptionv3[46] | 67.748% | 71.602% | 71.965% | 70.438% | 77% |
| se-resnet50[17] | 67.847% | 72.040% | 71.726% | 70.538% | 81% |
| DFL<br>(vgg16_bn)[39] | 67.519% | 69.511% | 71.180% | 69.403% | 75% |
| NTS<br>(resnet50)[47] | 66.077% | 70.152% | 69.679% | 68.969% | 75% |
| PC<br>(resnet50)[40] | 67.060% | 71.096% | 70.020% | 69.392% | 78% |
| MC-loss<br>(resnet50)[48] | 65.045% | 68.916% | 68.405% | 67.455% | 81% |
| B-CNN(vgg16)[37] | 61.422% | 62.226% | 61.903% | 61.850% | 71% |

**Extended Data Table 1 | Performance evaluation results.**
|  |  |  |  |
| --- | --- | --- | --- |
| SE-resnet50[17] | Valid 1 | Valid 2 | Valid 3 |
| Training image number | 31569 | 31655 | 31688 |
| Validation image number | 3051 | 2965 | 2932 |
| Genus accuracy (41) | 79.49% | 82.43% | 84.60% |
| Genus Average accuracy | 81.8% |  |  |
| Species Accuracy (113) | 67.847% | 72.040% | 71.726% |
| Species Average accuracy | 70.538% |  |  |

**Extended Data Table 2.** Expert performance on golden standard test set.

|  | Genus Accuracy | Species Accuracy | Average time cost (min) |
| --- | --- | --- | --- |
| Academic experts (14) | 66.5% (95%CI 60.5%, 70.8%) | 50.4% (95%CI 44.7%, 56.0%) | 5.4 (95%CI 4.9, 5.8) |
| Industry experts (7) | 45.29% (95%CI 33.0%, 57.6%) | 34.9% (95%CI 22.7%, 47%) | 4.1 (95%CI 3.0, 5.2) |

**Extended Data Table 3.**
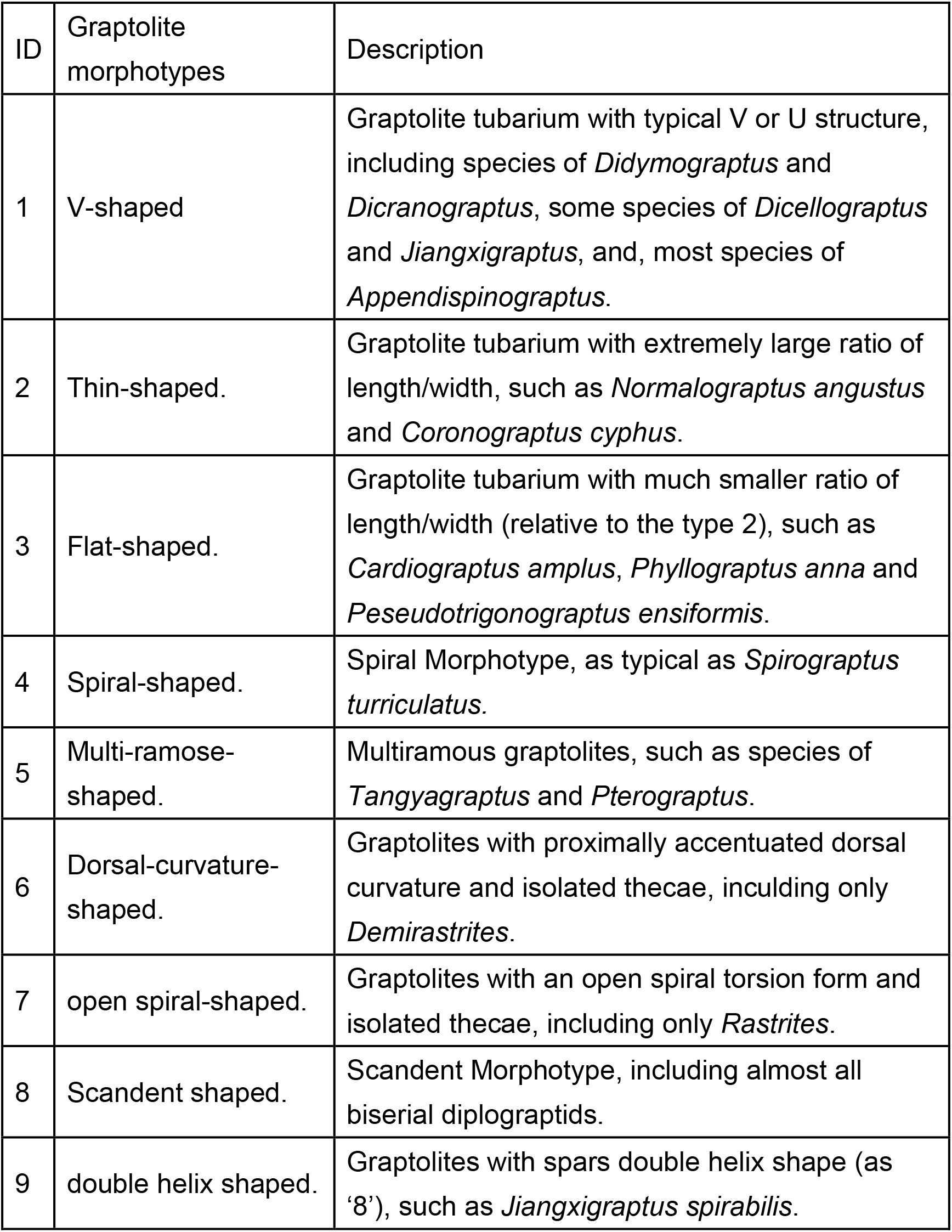
Nine graptolite morphotypes [33], clustered from our data set using UMAP.

## Appendix 1. A spreadsheet file showing our data source and data volume.

Fields include taxonomical names, revised names, specimen serial number, geological ages, locality, horizon, photo numbers, and related references of every specimens. Totally 1,565 pieces of specimens were used and all of them are deposited at the Nanjing Institute of Geology and Palaeontology (NIGP), Chinese Academy of Sciences (CAS). Honghe Xu. (2021). Graptolite specimens for AI training to improve shale gas exploration (Version 2021-5-24) [Data set]. Zenodo. http://doi.org/10.5281/zenodo.4782770

## Appendix 2. Video demonstrate our graptolite image dataset and the interactive Uniform Manifold Approximation and Projection Visualisation of the image dataset.

## Appendix 3. The blanket questionnaire we sent to palaeontological taxonomists and oil/gas industry specialists.

